# The homology analysis of ACE2 gene and its distinct expression in laboratory and wild animals

**DOI:** 10.1101/2021.04.08.439088

**Authors:** Gang Wang, A-Mei Zhang, Binghui Wang, Jianhua Yin, Yue Feng, zulqarnain Baloch, Xueshan Xia

**Affiliations:** Faculty of Life Science and Technology & Yunnan Provincial Center for Molecular Medicine, Kunming University of Science and Technology, Kunming 650500, Yunnan, China

**Author notes:** corresponding authors: Prof. Xueshan Xia, Faculty of Life Science and Technology, Kunming University of Science and Technology, Kunming, Yunnan 650500, China., Prof. Dr. Zulqarnain Baloch, Faculty of Life Science and Technology, Kunming University of Science and Technology, Kunming, Yunnan 650500, China.

**Keywords:** ACE2, expression, species, SARS-CoV-2, tree shrew, wild bats

## Abstract

Angiotensin-converting enzyme-2 (ACE2) has been recognized as an entry receptor of severe acute respiratory syndrome coronavirus 2 (SARS-CoV-2) into the host cells while bats has been suspected as natural host of SARS-CoV-2. However, the detail of intermediate host or the route of transmission of SARS-CoV-2 is still unclear. In this study, we analyze the conservation of ACE2 gene in 11 laboratory and wild animals that live in close proximity either with Bats or human and further investigated its RNA and protein expression pattern in wild bats, mice and tree shrew. We verified that the wild-bats and mice were belonged to Hipposideros pomona and Rattus norvegicus, respectively. ACE2 gene is highly conserved among all 11 animals species at the DNA level. Phylogenetic analysis based on the ACE2 nucleotide sequences revealed that wild bat and Tree shrew were forming a cluster close to human. We further report that ACE2 RNA expression pattern is highly species-specific in different tissues of different animals. Most notably, we found that the expression pattern of ACE2 RNA and protein are very different in each animal species. In summary, our results suggested that ACE2 gene is highly conserved among all 11 animals species. However, different relative expression pattern of ACE2 RNA and protein in each animal species is interesting. Further research is needed to clarify the possible connection between different relative expression pattern of ACE2 RNA and protein in different laboratory and wild animal species and the susceptibility to SARS-CoV-2 infection.

## Introduction

Coronavirus disease 2019 (COVID-19) pandemic, which is caused by the novel severe acute respiratory syndrome coronavirus 2 (SARS-CoV-2), has become a severe public health threat in the world [1]. As of February 1, 2021, SARS-CoV-2 has infected 102584351 peoples and resulting in more than 2222647 deaths, according to data from the World Health Organization (WHO)[2]. SARS-CoV-2 is a positive-sense, enveloped, single-stranded RNA virus, with 2 open reading frames (ORFs), with approximately 30 kb genome, and belonging to the genus Betacoronavirus. SARS-CoV-2 is 80~120nm in diameter, spherical in shape, nucleic acid and nucleocapsid protein are wrapped by lipid bilayer membranes. There are three important structural proteins: S, E and M proteins on the surface of the lipid bilayer [3]. Seven strains of SARS-CoV-2 have been reported so far and comparison of whole genome sequence of human CoVs strains has suggested that all strains are identical [4–6]. Moreover, a comparison of SARSCoV-2 genome with other coronavirus sequences shows that SARS-CoVs are probably originated in bats and then spread to intermediate hosts, before infecting humans. Therefore, these viruses may also have a wider host range [7, 8].

A comparative genetic sequence analysis of the SARS-CoV-2 and SARS-CoV sequences showed that the whole spike protein, receptor-binding domain (RBD), and receptor binding motif (RBM) sequence similarity was 76% to 78%, 73% to 76%, and 50% to 53%, respectively. This high sequence identity suggested that SARS-CoV-2 may have same entry receptor Human ACE2. ACE2 gene was discovered in 2000 and consists of 805 amino acids. ACE2 is widely expressed in ileum, duodenum, jejunum, colon in kidney, testis, heart and gastrointestinal system throughout the animal kingdom from fish, amphibians, reptiles, birds to mammals [9–11]. Its main function is to regulate and maintain the relative stability of the body environment, balance of blood pressure and water and electrolytes. [12–14]. It has been showed that RBD of 2019-nCoV bind to the peptidase domain (PD) of ACE2 from different animal hosts. These observations provide important information on the SARS-CoV-2 interaction with ACE2. So, we can propose that SARS-CoV-2 may be transmitted from animals to humans through spillover or cross-species transmission. However, these results are not enough to confirm the source of SARS-CoV-2, to explain its intermediate host or route of transmission to humans. Additionally, keeping in view the different expression pattern of ACE2, the presence of structural and sequence variants in different animals may have effect on susceptibility and immune response to SARS-CoV-2. Therefore, this study was designed to analyze the conservation of ACE2 gene in 11 different laboratories and wild animals species that live in close proximity either with Bats or human and further investigated its RNA and protein expression pattern in wild bats, mice and tree shrew (Tupaia belangeri).

## Materials and Methods

### Ethical statement

In this study, all the animal related to experiments were strictly accomplished following the instruction of Chinese Regulations of Laboratory Animals (Ministry of Science and Technology of the People’s Republic of China) and Laboratory Animal—Requirements of Environment and Housing Facilities (GB 14925-2010, National Laboratory Animal Standardization Technical Committee). All animal experiments were performed under sodium pentobarbital anesthesia. This study was approved by the Animal Experiment Committee in Kunming University of Science and Technology China.

### Experimental sample

Wild bats (Hipposideros Pomona) and mice (Rattus norvegicus) were caught from the wild forest in PuEr and Kunming of Yunnan Province respectively. Oryctolagus cuniculus f. domesticus, Fancy Rat, Golden hamster, Sprague-Dawley Rat and Phodopus sungorus were purchased from local pet animal market. Mesocricetus auratus, tree shrew, and C57 were commercially provided by the Laboratory Animal Resource Center of Kunming Medical University. All animals were anesthetized with thiopental sodium (80 mg/kg), heart, liver, spleen, lung, kidney, brain, colon, duodenum, stomach and trachea tissues were dissected and preserved in liquid nitrogen till further use.

### β-globin gene sequencing and identify of wild animal species

50mg kidney tissues of Bat and Mice were individually homogenized thoroughly. The genomic DNA extracted by TIANamp Genomic DNA Kit (Lot#U8923), according to the manufacture instructions, beta-globin genes were amplified by PCR to determine DNA stability and the DNA was preserved at −20 °C. The quality of DNA of all samples was tested for β-globin. The DNA extracts were used as a template for PCR amplification using species-specific primers. The PCR products was directly sent to TSINGKE Biological Technology for sequencing using the ABI PRISM Big Dye Terminator Cycle Sequencing Ready Reaction (Invitrogen, Beijing, CN) on an ABI 310 DNA analyzer. After the sequence was obtained, confirmed to be β-globin gene sequence by BLST, in NCBI database. Then the β-globin gene sequences of different bats and mice were downloaded from the database, and the species names of wild bats and mice were further determined by constructing a phylogenetic tree.

### ACE2 gene sequencing and analysis

The RNA of the heart, liver, spleen, lung, kidney, brain, colon, duodenum, stomach and trachea of the 11 species were extracted using RNAiso Plus (TaKaRa, Lot#AJF1819A), and reverse-transcribed using TaKaRa PrimerScript ^TM^ II 1st Strand cDNA Synthesis Kit (Lot#AK70993A). The PCR products were commercial sequenced as described above. Multi-sequence alignment of the ACE2 of the 11 species was performed using ClustalW, the gene sequence comparison and Neighbor-Joining tree were performed with MEGA 7.0 software [15]. The bootstrap value was 1000 and the ACE2 sequence of human was downloaded from GeneBank (accession number: NM_001371415.1). The translation of ACE2 gene sequences of 11 species into amino acid sequences was also completed by MEGA7.0 software, and then we compared the differences in the key sites of the binding of ACE2 protein and SARS-CoV-2Spike protein in these 11 species, which were reported by Fang S et al[15]. The conservation of ACE2 key site residues is accomplished by WebLogo[16].

### Determine on expression of ACE2 in tissues

For determining ACE2 gene expression in different tissues, the relevant reference sequences were downloaded from the public database of NCBI, and primers were designed with PRIMER5 (Table 2). The reverse transcribed cDNA was used as a template for qPCR amplification. QPCR was performed using Power SYBR Green PCR Master Mix (Vazyme ChamQ SYBR qPCR Master Mix,vazyme code:Q311-02)) in Biosystems ABI 7500 machine.

**Table 1:**
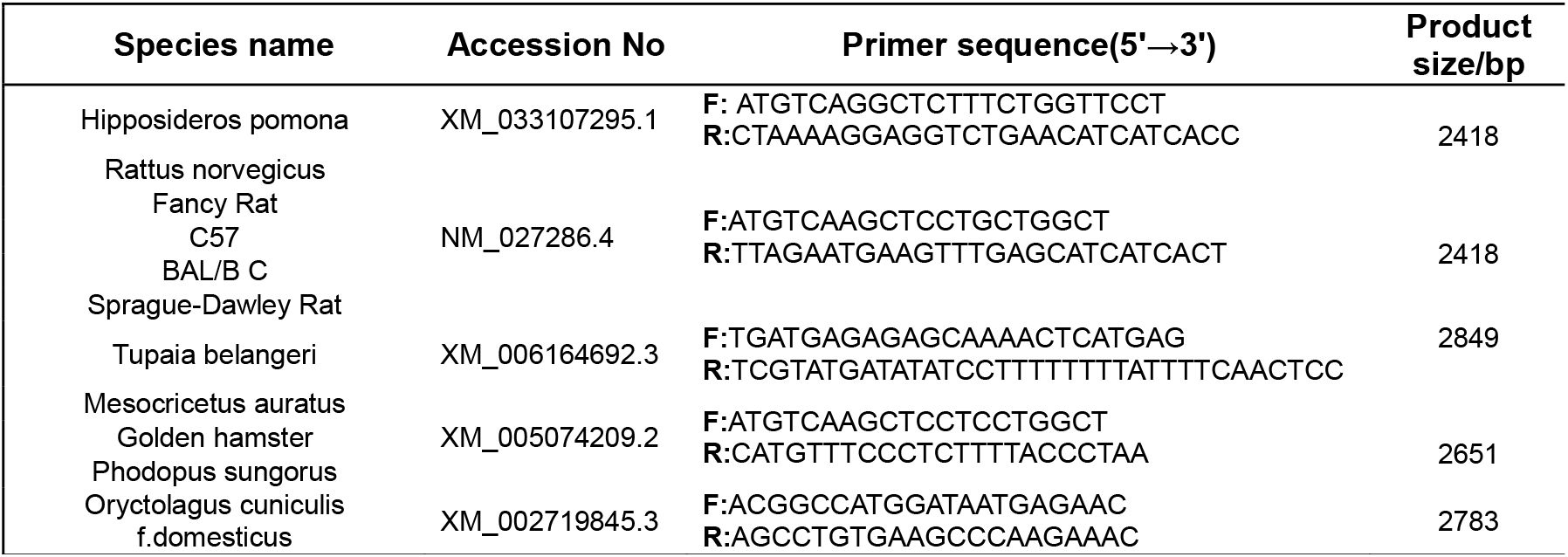
ACE2 gene primer sequence

**Table 2:** Sequence alignment of 14 key polar residues in 11different species

| Species name |  | Key polar residues of ACE2 in different species |  |  |  |  |  |  |  |  |  |  |  |  |  |  |  |  | Sequence similarity |  |  |
| --- | --- | --- | --- | --- | --- | --- | --- | --- | --- | --- | --- | --- | --- | --- | --- | --- | --- | --- | --- | --- | --- |
|  |  | 24 | 27 | 28 | 30 | 31 | 34 | 35 | 37 | 38 | 41 | 42 | 79 | 82 | 83 | 330 | 353 | 355 | 357 | Full | Key polar |
| Homo sapiens |  | Q | T | F | D | K | H | E | E | D | Y | Q | L | M | Y | N | K | D | R | 100% | 100% |
|  | Wild bat | L | T | F | E | K | S | E | E | D | Y | Q | L | N | Y | N | K | D | R | 79.8% | 78.57% |
| Wild mice |  |  |  |  |  |  |  |  |  |  |  |  |  |  |  |  |  |  |  | 81.0% | 57.14% |
|  |  | D | V | F | D | E | I | E | E | E | Y | Q | Q | R | Y | K | N | D | R |  |  |
| Fancy Rat |  |  |  |  |  |  |  |  |  |  |  |  |  |  |  |  |  |  |  | 81.9% | 57.14% |
|  |  | N | T | F | N | N | Q | E | E | D | Y | Q | T | S | F | N | H | D | R |  |  |
| Rattus norvegicus |  |  |  |  |  |  |  |  |  |  |  |  |  |  |  |  |  |  |  | 82.1% | 64.29% |
|  |  | S | S | F | N | K | Q | E | E | D | Y | Q | I | N | F | N | H | D | R |  |  |
| C57 |  |  |  |  |  |  |  |  |  |  |  |  |  |  |  |  |  |  |  | 81.7% | 57.14% |
|  |  | N | T | F | N | N | Q | E | E | D | Y | Q | T | S | F | N | H | D | R |  |  |
| BAL B/C |  |  |  |  |  |  |  |  |  |  |  |  |  |  |  |  |  |  |  | 82.1% | 57.14% |
|  |  | N | T | F | N | N | Q | E | E | D | Y | Q | T | S | F | N | H | D | R |  |  |
| Sprague-Dawley Rat |  |  |  |  |  |  |  |  |  |  |  |  |  |  |  |  |  |  |  | 82.1% | 64.29% |
|  |  | K | S | F | N | K | Q | E | E | D | Y | Q | I | N | F | N | H | D | R |  |  |
| Oryctolagus cuniculus f. domesticus |  |  |  |  |  |  |  |  |  |  |  |  |  |  |  |  |  |  |  | 85.3% | 78.57% |
|  |  | L | T | F | E | K | Q | E | E | D | Y | Q | L | T | Y | N | K | D | R |  |  |
| Mesocricetus auratus |  |  |  |  |  |  |  |  |  |  |  |  |  |  |  |  |  |  |  | 84.6% | 92.86% |
|  |  | Q | T | F | D | K | Q | E | E | D | Y | Q | L | N | Y | N | K | D | R |  |  |
| Golden hamster |  |  |  |  |  |  |  |  |  |  |  |  |  |  |  |  |  |  |  | 84.5% | 92.86% |
|  |  | Q | T | F | D | K | Q | E | E | D | Y | Q | L | N | Y | N | K | D | R |  |  |
| Phodopus sungorus |  |  |  |  |  |  |  |  |  |  |  |  |  |  |  |  |  |  |  | 83.0% | 92.86% |
|  |  | Q | T | F | D | K | Q | E | E | D | Y | Q | L | N | Y | N | K | D | R |  |  |

Nonpolar residues: 27, 28, 79 and 82; 14 polar residues: 24, 30, 31, 34, 35, 37, 41, 42, 83, 330, 353, 355 and 357.

Protein extracts were generated from tissues by using RIPA buffer. The protein concentration of each tissue was determined by BCA method (TaKaRa, Lot#A4401A). The tissue protein was used for SDS-PAGE electrophoresis at the same quality, and then transferred to the PVDF membrane by the wet spinning method. ACE2 protein primary antibody (BOSTER, Lot#BST19874624) and β-actin protein primary antibody (ABclonal, Lot#9100026001) respectively, and incubated overnight at 4 °C. After washing the membrane with PBST, the second antibody of IgG (HRP) was transferred (ABCOM, Lot#GR 32999244-7). After 2 hours of incubation at room temperature, membrane was photographed in the automatic imaging luminescence system.

## Result

### Species identification of wild bats and mice

Wild bats and mice are widely populated together in the forest of PuEr and Kunming. Wild bats have already been suspected as a possible host of SARS-CoV and SARS-CoV-2. While, wild mice are living in the same environment from millions of years together with wild bats. Therefore, It can be suspected that there may be an interaction between wild bats and wild mice in transmission of SARS-CoV-2. At first, we were not sure about the species of wild bats and mice. Therefore, we first examined the mitochondrial loci cytochrome b (cyt b) for differentiating wild bats and mice species by using the primers reported by Irwin[17]. Results showed that the wild-bats were belonged to Hipposideros pomona and the wild mice were Rattus norvegicus (**Figure 1**).

**Figure 1:**
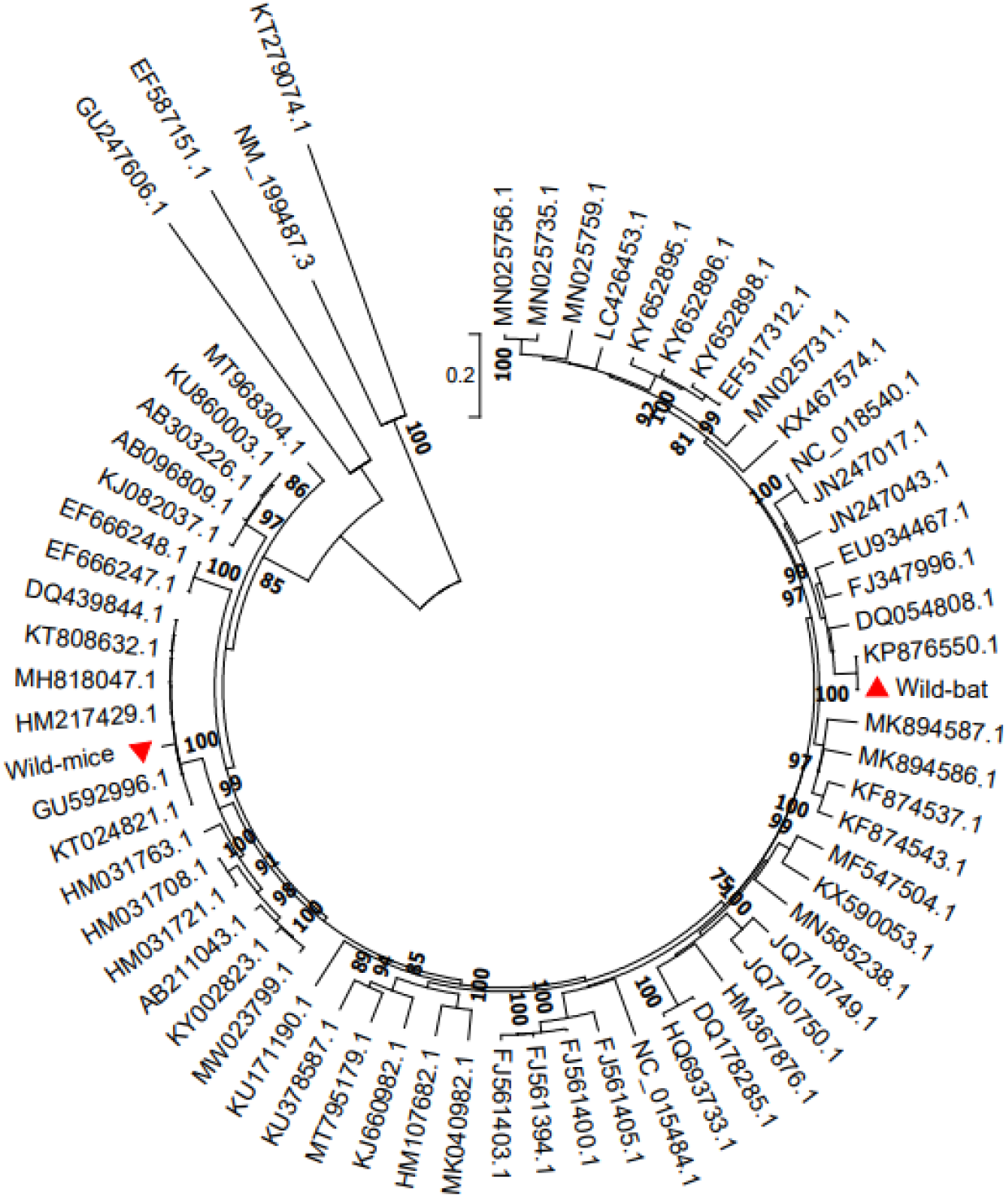
Species identification of wild bat and mice(Neighbor-Joining tree)

### Analysis of ACE2 nucleotide homology

The ACE2 genes of these 11 species amplified by PCR with designed primers (**Table 1**). We found that ACE2 gene is highly conserved among selected 11 animal’s species at the DNA level. Overall, the nucleotide homology between Oryctolagus cuniculis f.domesticus and human ACE2 is 88.2%, Oryctolagus cuniculis f.domesticuss and human is 87.2%, wild bat and human ACE2 is 84.8%, wild mice and human ACE2 is 84.9%. While, the nucleotide sequence homology of human ACE2 with other species is between 85.1% and 87.1% (**Figure 2A**).

**Figure 2:**
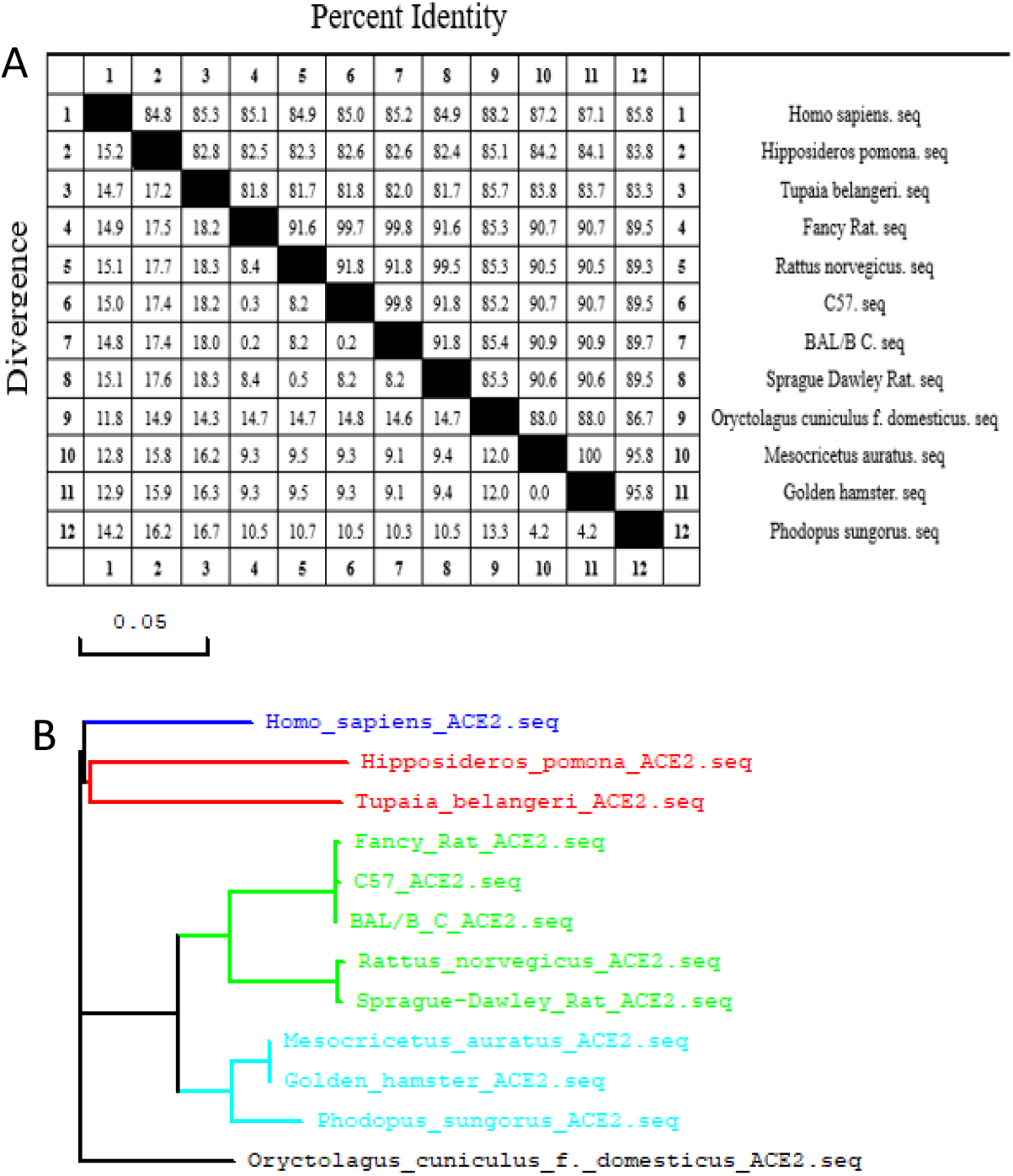
Homology analysis and biological phylogenetic tree of ACE2. A. Homology analysis of ACE2 nucleotide. **B.** Biological phylogenetic tree of ACE2 gene among different species.

Phylogenetic analysis of the complete ACE2 nucleotide sequences of 11 species revealed that wild bat and Tree shrew were forming a cluster close to human. While, wild mice, C57 and BALB/C were fall in same branch and suggests a close genetic relationship between them. Wild mice and Sprague-Dawley Rat were formed a tight cluster in the distinct branch; Mesocricetus auratus and Golden hamster were grouped together (**Figure 2B).** Oryctolagus cuniculis f.domesticus was clustered into a distinct clade.

### Key polar residues of ACE2 binding to SARS-CoV-2 RBD

The nucleotide sequences of 11 animal species ACE2 gene were translated into amino acid sequences (**Table 2)**. In this study, we found total 18 residues on ACE2 from all the 11 species which have interactions with 2019-nCoV-Spike. Among them, 4 were non-polar residues and 14 were polar-residues. We found that there are total 9 (28, 35, 37, 38, 41, 42, 330, 355 and 357) highly conserved residues which may have key role in the basic binding affinity with the novel coronavirus. Similarly, a total of 5 un-conserved residues were found which may have a decisive effect on the binding affinity with the novel coronavirus (**Figure 3**).

**Figure 3:**
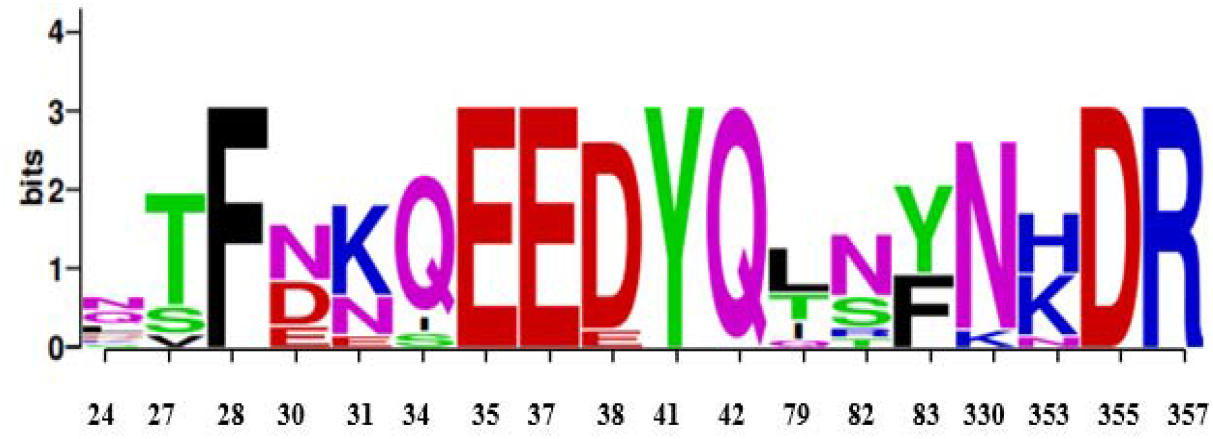
Sequence alignment result of 18 key residues of ACE2 protein in 11 species that form the interactions with SARS-CoV-2-Spike. Nonpolar residues: 27,28,79 and 82; 14 polar residues: 24,30,31,34,35,37,41,42,83,330,353,355 and 357.

By sequence alignment and statistical analysis, the sequence similarities of ACE2 proteins for all 11 species were higher than 79% compared to human **(Table 2).** Compared with the human ACE2 amino acid sequence, the wild bats ACE2 amino acid sequence had the lowest homology (79.8%); and Oryctolagus cuniculis f.domesticus has the highest homology (85.3%). Further we found that the sequences of 14 key polar amino acids forming the binding interface from different species were quite different, especially for Fancy rat, C57, tree shrew and BAL B/C, which has 57.14% similarity with human beings. Next, we calculated the binding free energy of each ACE2/SARS-CoV-2S protein complex and checked the stability of each ACE2/SARSCoV-2S protein complex.

### Expression of ACE2 gene in different tissues of wild bat, tree shrew and wild mice

In this study, phylogenetic tree generated using genomic data; wild bat and tree shrew were selected for further analysis, because the clustered of both species are close to humans. Additionally, as wild mice are widely populated together with Bats, thus we also added wild mice for further analysis. RNA was extracted from heart, liver, spleen, lung, kidney, brain, colon, duodenum, stomach and trachea tissues of wild bats, mice, and tree shrew and RT-qPCR was performed using specific primers (**Table 3**). The relative mRNA level of ACE2 in different tissues of wild bats, mice, and tree shrew has been shown in **Fig 4.** The mRNA of ACE2 was high in stomach of wild bats followed by kidney, liver, colon, duodenum, trachea, heart, spleen, and brain (**Fig 4A)**. The mRNA of ACE2 was high in stomach of wild mice followed by duodenum, colon, kidney, trachea, heart, lung, brain, liver, and spleen (**Fig 4B).** Similarly, the mRNA of ACE2 was high in colon of tree shrew followed by duodenum, kidney, heart, stomach, trachea, brain, liver, spleen (**Fig 4C)**. Surprisingly, there was no expression of ACE2 gene in lungs of wild bats, mice, and tree shrew.

**Figure 4:**
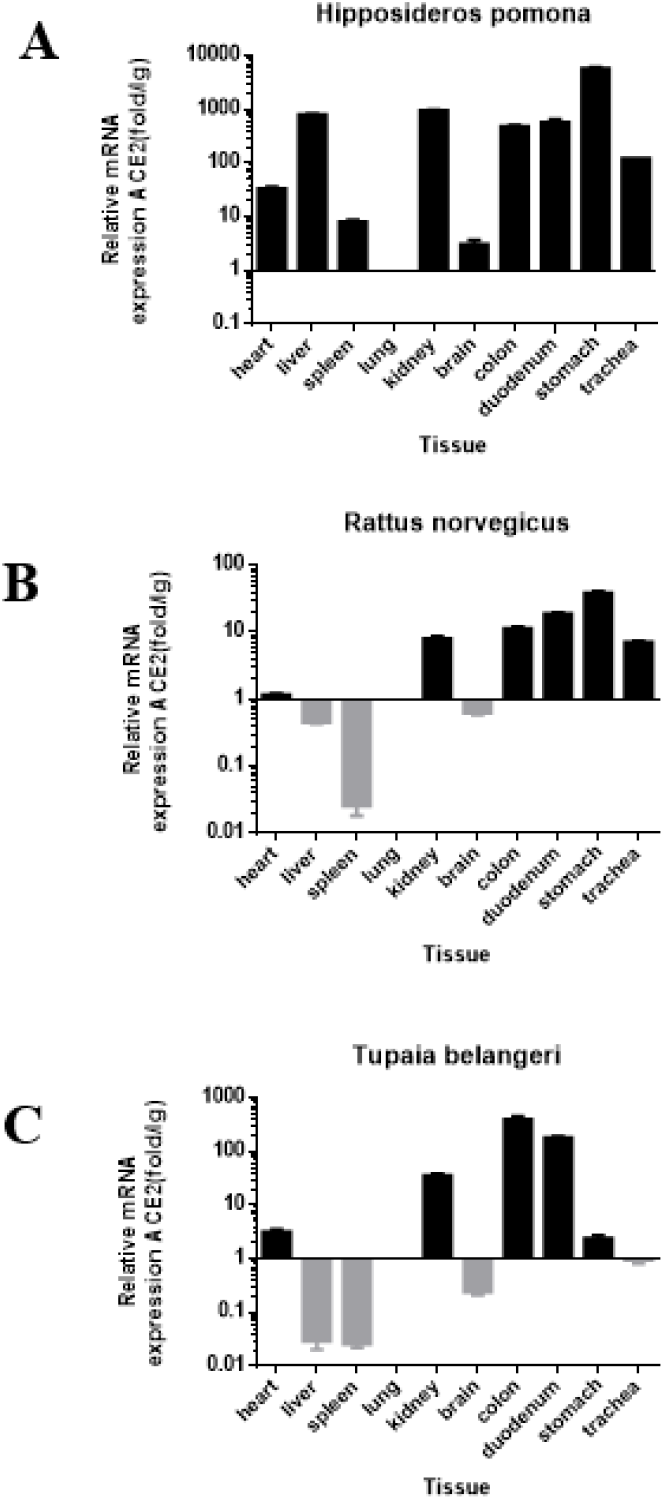
Expression pattern of ACE2 RNA in organs of Hipposideros pomona, Rattus norvegicus and Tupaia belangeri tissues.

**Table 3:**
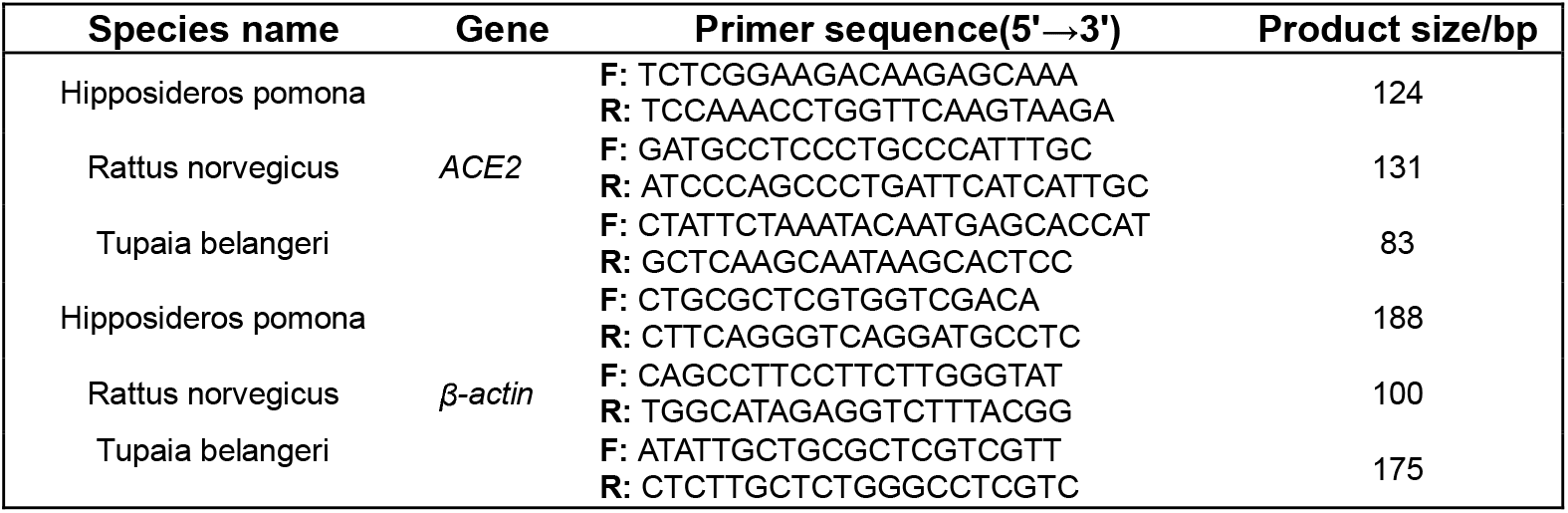
ACE2 gene Real-Time PCR primer sequence

Several recent investigations have revealed the mRNA of ACE2 in human tissues but nobody yet explored ACE2 protein expression in different mammalian species. We profiled and normalized the expression patterns of ACE2 protein in different organs of wild bat, mice, and tree shrew (**Fig 5).** The level of ACE2 protein was high in the liver of wild bat followed by kidney, colon, duodenum, stomach and trachea, suggesting that the liver, kidney, colon, duodenum, stomach and trachea could be highly vulnerable to SARS-CoV-2 infection in wild bats **(Fig 5A).** The expression of protein was high in kidney of wild mice followed by colon, duodenum, stomach and trachea, suggesting that the kidney, colon, duodenum, stomach and trachea could be highly vulnerable to SARS-CoV-2 infection in wild mice (**Fig 5B)**. The expression of ACE2 protein in tree shrew has been shown in **fig 5C**. Results showed that colon, duodenum and kidney could be highly vulnerable to SARS-CoV-2 infection in tree shrew.

**Figure 5:**
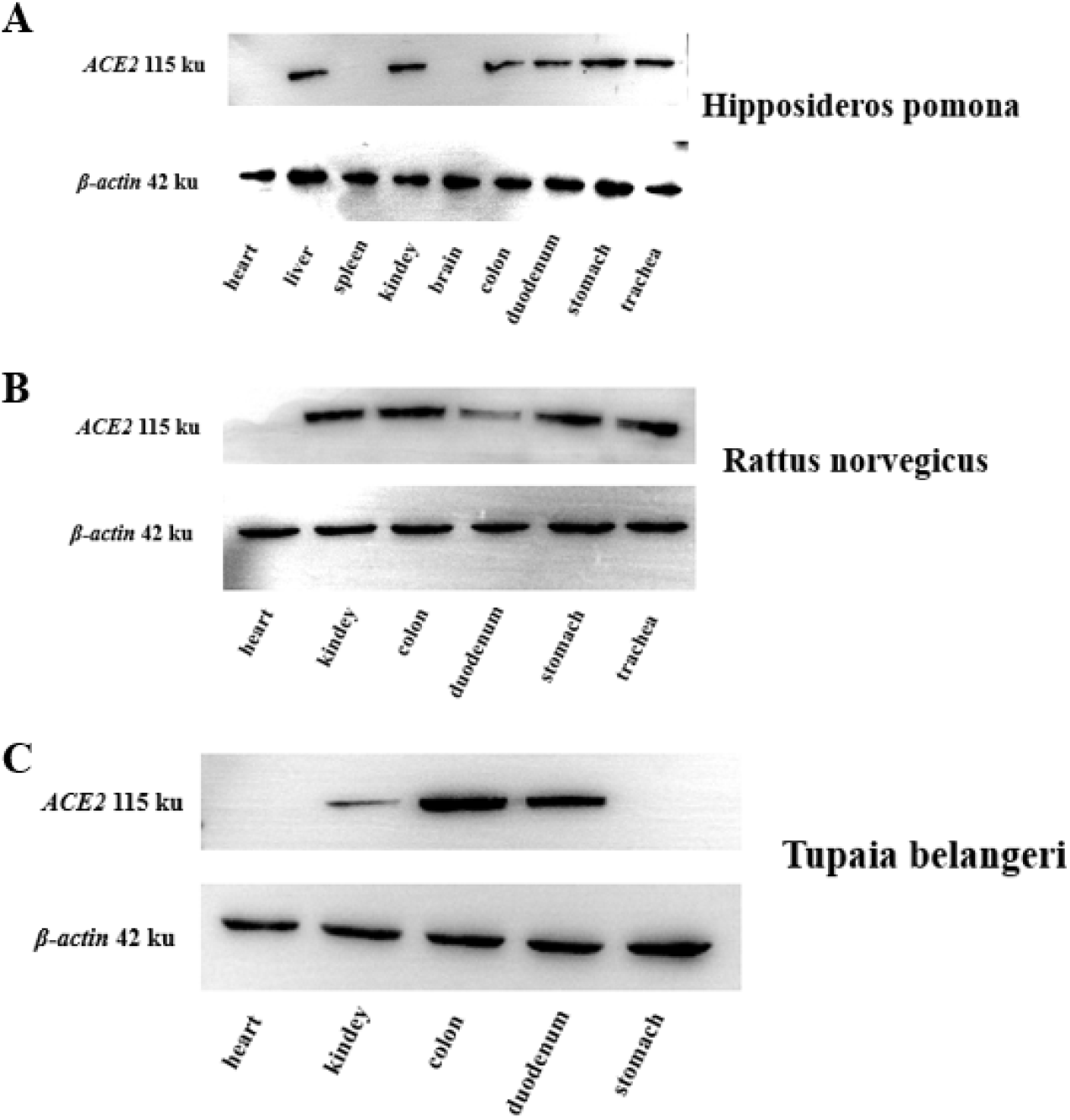
Expression pattern of ACE2 Protein in different organs of Hipposideros pomona, Rattus norvegicus and Tupaia belangeri

## Discussion

The research on ACE2 was initially involved in the humoral regulation of cardiovascular system as an important member of renin-angiotensin system(RAS), but in subsequent studies, ACE2 has been proved to be the receptor of SARS-CoV, SARS-CoV-2 and HCoV-NL63 coronaviruses[7, 18, 19]. Further studies suggested that ACE2 Key polar residues played key roles in the binding and maintaining the stabilities of ACE2/2019-nCoV-Spike RBD interactions across multiple mammalian orthologs. However, these studies only used public database sequences without any laboratory based experiments. In this study, we have collected samples from wild, pet and laboratory animals to characterize the conservation of ACE2 in different animals and investigated the expression patterns of ACE2 gene in different mammalian species.

After declaring ACE2, as a possible receptor of SARS-CoV-2, a wide range of animal species has been investigated in this and previous studies [20, 21]. Our results show that overall ACE2 sequences are highly conserved across all recruited animals’ species which is in line with previous reports [21, 22]. This conservation shows that SARS-CoV-2 may have a broad range of animal hosts [22]. Most notably, in the phylogenetic tree based on the ACE2 nucleotide sequences shows that the wild bats and tree shrew are the species clustered closest to humans. This highlights a close evolutionary relationship between wild bats (and tree shrew. As tree shrew have already been successfully developed as animal models of different viral infections including influenza [23], herpes simplex [24], hepatitis C virus [25], and ZIKV [26]. We can hope may be tree shrew will also become SARS-CoV-2 infection animal model.

We further investigated the interaction between ACE2 and SARS-CoVS protein, our results shows that there are total 18 key amino acids on ACE2 that can bind to the binding domain of S protein receptor [27]. Among these 18 key amino acids, 14 amino acids belong to polar interaction (hydrogen bond, salt bridge, etc.), which plays an absolutely major role in maintaining the stability of ACE2/SARS-CoV-2 S protein interaction[15]. Further, we compared the full length amino acid sequences of 11 species with humans and the 14 key polar amino acids. It was found that the total length of amino acids was not directly related to 14 key polar amino acids and sequences of 14 key polar amino acids forming the binding interface from different species were quite different among recruited species. In this study, the similarity of 14 key polar amino acids was the highest among Mesocricetus auratus, Golden hamster and Phodopus sungorus. It is consistent with the reported Mesocricetus auratus which is expected to become a model of SARS-CoV-2 infection [28]. Is there any possible risk of SARS-CoV-2 infection in pet hamsters (Golden hamster, Phodopus sungorus) that are in close contact with humans?

It is well accepted that expression of ACE2 RNA and protein plays an essential role in mediating SARS-CoV-2 infection in the host cells[22, 29], however, the expression of ACE2 is highly tissue specificity [9, 10]. In this study, we determine the relative expression of ACE2 RNA and protein in different tissues of wild bats, mice and tree shrew. Previously reported studies, mostly surveyed on the expression of ACE2 RNA without considering the importance of protein data[22]. While some human based studies had established that the expression of ACE2 RNA and protein are almost similar [29, 30] which are inconsistant with our results (**Figure 4** and **Figure 5**).

## Conclusion

The results of this study suggested that ACE2 gene is highly conserved among all 11 laboratory and wild animals species. However, different relative expression pattern of ACE2 RNA and protein in each animal species is interesting. Further research is needed to clarify the possible relationaship between different relative expression pattern of ACE2 RNA and protein in different laboratory and wild animals’ species and the susceptibility to SARS-CoV-2 infection.

## Author contribution

Xia Xueshan, designed and supervised the study, Gang Wang, A-Mei Zhang, Binghui Wang, Jianhua Yin, and Yue Feng finished the lab work, bioinformatic process as well as statistical analysis, Xia Xueshan and Zulqarnain Baloch were responsible for manuscript writing and revising.

## Figure legends

**Figure 1**: Species identification of wild bat and mice (Neighbor-Joining tree).

**Figure 2**: Homology analysis and Neighbor-Joining tree of ACE2.

A. Homology analysis of ACE2 nucleotide. B. Neighbor-Joining tree of ACE2 gene among different species.

**Figure 3**: Sequence alignment result of 18 key residues of ACE2 protein in 11 species that form the interactions with SARS-CoV-2-Spike. Nonpolar residues: 27,28,79 and 82; 14 polar residues: 24,30,31,34,35,37,41,42,83,330,353,355 and 357.

**Figure 4**: Expression pattern of ACE2 RNA in organs of wild bat, wild mice and Tupaia belangeri tissues.

**Figure 5**: Expression pattern of ACE2 Protein in different organs of wild bat, wild mice and Tupaia belangeri.

